# Dual-Route H5N1 Vaccination Induces Systemic and Mucosal Immunity in Murine and Bovine Models

**DOI:** 10.1101/2025.09.21.677614

**Authors:** Joshua Wiggins, Adthakorn Madapong, Eric A Weaver

## Abstract

Since its discovery in U.S. dairy cattle in early 2024, the highly pathogenic H5N1 avian influenza (clade 2.3.4.4b) has spread widely among herds, causing major economic losses. This zoonotic event emphasizes the urgent need for H5 vaccines that elicit strong, durable, cross-reactive immune responses in cattle, especially young calves. To address this, we immunized mice and cattle with a centralized consensus H5 vaccine, designed to localize to the central node of the human H5 phylogenetic tree. The vaccine was delivered using serotype-switched adenoviral vectors in a prime–boost regimen, combined with intramuscular and intranasal coadministration to target systemic and mucosal immunity and elicit strong humoral and cellular immune responses. This approach strategically integrates multiple innovative features: centralized consensus immunogens, mucosal targeting, and vector serotype switching that are aimed at maximizing immune protection against H5N1 viruses. Our results show that vaccination elicited strong humoral and cellular immunity in both mice and calves. In challenge experiments, vaccinated mice were fully protected against lethal infection with multiple divergent H5N1 strains, including the 2024 bovine isolate (A/bovine/Ohio/B24OSU-439/2024). Given that vaccine induced immunity was consistent across species, these results support the translatability of the mouse model findings to cattle. Overall, our findings represent a promising approach for immunizing cattle and other key livestock against HPAI H5N1, mitigating agricultural losses, and reducing the risk of zoonotic transmission.

**Significance Statement:** H5N1 influenza A virus is a serious pathogen recently detected in many mammals, including cattle. It transmits sporadically to humans and causes major economic losses in dairy and poultry industries, raising global concern. No H5N1 vaccines are currently licensed for cattle. To address this gap, we tested a centralized consensus H5 (H5CC) vaccine, previously effective in mice and swine. Using multiple Adenovirus vectors, we delivered H5CC intramuscularly and intranasally to one-week-old calves. The vaccines induced strong humoral and cellular immune responses, essential for preventing infection and limiting transmission. These findings highlight the vaccine’s potential to reduce agricultural losses and remove cattle as a newly established reservoir for zoonotic spread, providing a promising strategy for mitigating pandemic risk.

## Introduction

Highly pathogenic avian influenza (HPAI) H5 influenza A virus (IAV) emerged in Guangdong, China in 1996 (A/goose/Guangdong/1/96) and has diversified into multiple clades, including the globally dominant clade 2.3.4.4b H5 lineage (1). This clade has caused widespread avian outbreaks leading to severe economic losses in the poultry industry and has demonstrated zoonotic transmission to humans, swine, marine mammals, companion animals, and notably dairy cattle (2-5). In March 2024, HPAI H5N1 genotype B3.13 (clade 2.3.4.4b) was first detected in U.S. dairy cattle (6). By mid-2025, it had been confirmed in 18 states across 1080 herds (7). Viral RNA and antigen have been detected in unpasteurized milk, mammary tissue, and nasal secretions of infected dairy cows (6, 8-10). Affected cows lose ∼900 kg of milk over a 60-day period, and economic losses from milk loss, mortality, and culling costs are estimated at ∼$950 per clinically affected cow (11). Sporadic zoonotic spillovers to humans have occurred, typically causing mild disease, though one fatality was reported in January 2025 (12, 13). The expanding mammalian host range and rising incidence of zoonotic transmission increases risks of mammalian adaptation and potential human-to-human spread (14). Bovine H5 vaccination could mitigate spread and economic impact. To date, a few experimental H5 vaccines have been tested in cattle (15-17), but no licensed vaccines are currently available.

Adenovirus (Ad) vectors are a highly attractive influenza vaccine delivery platform due to their capacity to induce durable, broad-spectrum immunity, eliciting potent humoral and cellular responses (18-20). Ad vectors can be engineered to be replication-defective via deletion of the E1 region, or as replication-competent by retaining E1 and deleting non-essential genes such as E3. These design decisions influence safety, antigen expression, and immunogenic persistence (18). Ad vectors also exhibit broad cellular tropism, targeting both mucosal epithelial cells and antigen-presenting dendritic cells, thereby facilitating intranasal and intramuscular vaccine administration (21, 22). Our lab has previously assessed the potential of Ad-vectored H5 influenza vaccines administered either intramuscularly or intranasally in mice and swine (23-25). Here, we test a H5 centralized consensus (H5CC) immunogen that phylogenetically localizes near the ancestral A/goose/Guangdong/1/96 (H5N1) virus strain. This immunogen was inserted into either replication-defective, high-seroprevalent species C adenoviruses (Ad5 and Ad6), or replication-competent, low-seroprevalent species D adenoviruses (Ad28 and Ad48). The vaccines were evaluated for immunogenicity through intramuscular and intranasal coadministration in murine and bovine models. We evaluate mucosal, humoral, and cell-mediated immune responses in both animal models and further assess cross-clade protection from lethal modified HPAI H5N1 viral challenges in mice.

## Results

### Development and characterization of Ad_H5CC vaccines

We developed a broadly protective vaccine using a consensus-based approach aimed at targeting H5Nx influenza, positioning the immunogen near the center of the H5 phylogenetic tree (Figure 1A). To assess the genetic diversity of H5 hemagglutinin (HA) strains, we analyzed 4,609 unique global H5 HA sequences from the GISAID database using Geneious Tree Builder (v11.1.5). A neighbor-joining phylogenetic tree was constructed with the Jukes-Cantor distance model and UPGMA method. As shown in Figure 1B, the recent evolutionary trajectory (blue nodes), reveals a significant expansion of strains within the 2.3.4.4b clade in 2022 when compared to 2020, suggesting heightened selection pressure likely driven by adaptation to mammalian hosts. Strains linked to transmission and disease in dairy cattle, isolated in 2024, formed a distinct subclade within the 2.3.4.4b lineage (purple and red branches), contrasting with the broader phylogenetic tree that reflects over 60 years of viral evolution prior to 2020. Taken together, these findings highlight the capacity of H5 viruses for rapid evolution in response to intensified host-pathogen interactions.

**Figure 1:**
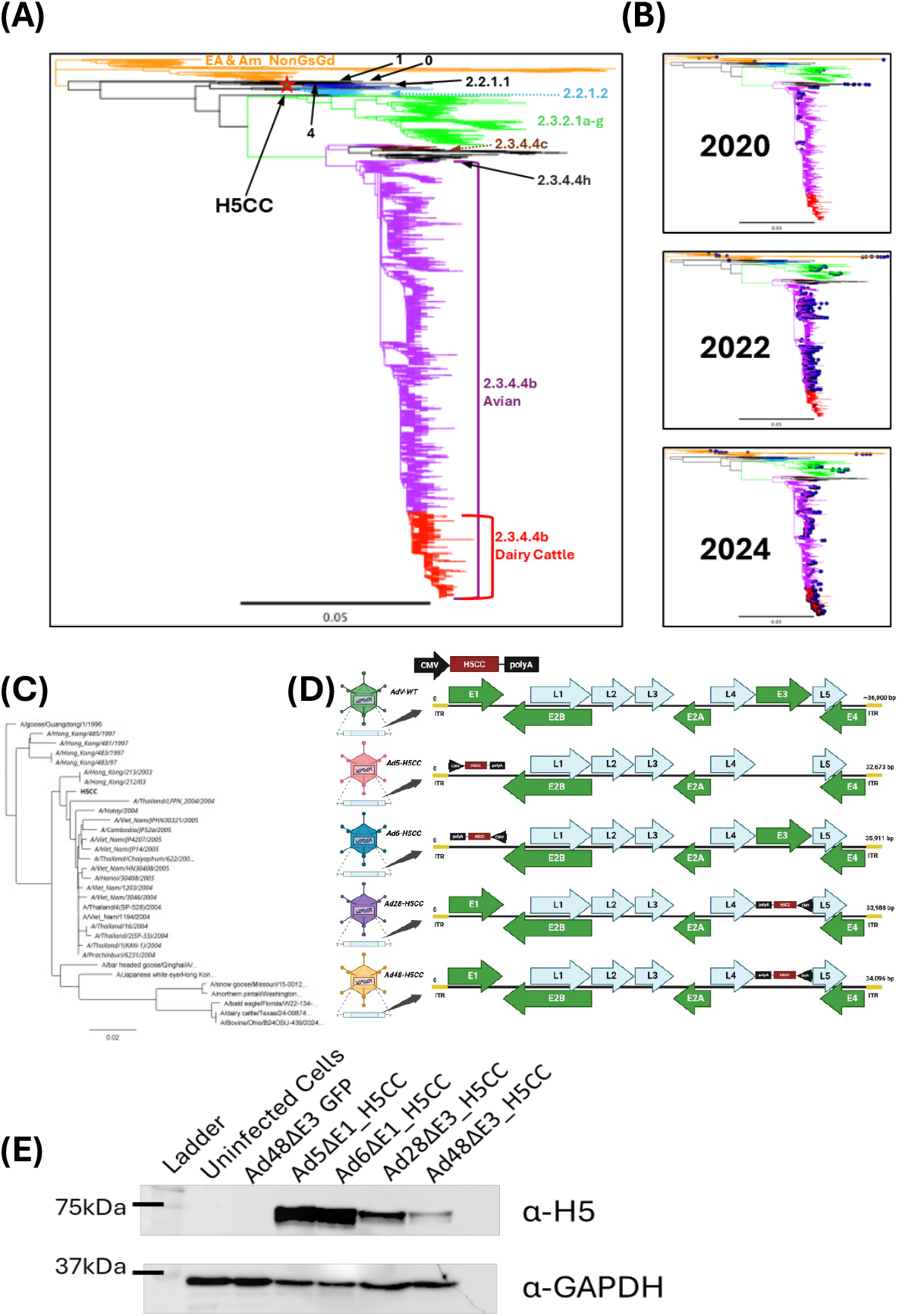
Phylogenetics of H5Nx; and H5CC vaccine design and characterization. (A); Genetic relationship between the vaccine candidate immunogen, H5CC (red star), and defined clades including ancestral Eurasian and American Non-Goose Guangdong (EA&Am NonGsGd) strains. Different colors depict various sub-clades, indicated to the right of the image. H5CC localizes to clade 1 and is closely related to A/Vietnam/1194/2004. (B); Evolution and genetic distribution of H5N1 viruses isolated in 2020, 2022, and 2024. Circulating strains for each year are marked with blue nodes. (C); Neighbor end-joining tree depicting the genetic relationship of all H5 strains used in this study. The 21 strains used to design the vaccine are shown in *italics* while the H5CC vaccine is shown in bold. (D); DNA schematic of a wild-type Adenovirus (AdV-WT) and the four vaccine vectors used in the study, including the E1 (replication deficient), or E3 (replication competent) deletions used to accommodate the vaccine immunogen. The size of each recombinant vector, as well as the orientation of the H5CC within the viral genome are depicted. (E); Western blot showing protein expression of the vaccine constructs in HEK 293 cells.

The synthetic H5CC vaccine was designed by aligning 21 unique clade 1 human H5 sequences from 1997 to 2005. The strains used to create the H5CC gene are shown in italics in Figure 1C, with accession numbers listed in the Materials and Methods section. Supplemental Figure 1 shows the H5CC vaccine is closely related to the clade 1 A/Vietnam/1194/04 (98.2% identity) and clade 0 A/goose/Guangdong/1/96 (96.0% identity) strains. Notably, the H5CC vaccine shares 91.7% identity with the currently circulating clade 2.3.4.4b bovine H5 IAV, despite being designed nearly two decades earlier and lacking a multi-basic cleavage site (Figures 1C and S1). To facilitate vaccine delivery, the H5CC construct was codon-optimized and cloned into four human adenovirus (Ad) vectors: Ad-5 and Ad-6 (high-seroprevalent species C), and Ad-28 and Ad-48 (low-seroprevalent species D). The H5CC gene was inserted under control of the strong cytomegalovirus (CMV) immediate-early promoter with a BGH-polyadenylation (PolyA) signal for enhanced stability. In the Ad-5 and Ad-6 vectors, the H5CC gene was cloned into the E1 region, rendering the vectors replication-deficient, while in the Ad-28 and Ad-48 vectors, the gene was inserted into the E3 region, making them replication-competent. Constructs were designated Ad5_H5CC, Ad6_H5CC, Ad28_H5CC, and Ad48_H5CC. Figure 1D depicts the vectors with their respective genome deletions and synthetic H5CC insertions. The vectors were verified by Oxford Nanopore sequencing to confirm their sequences, and H5 gene expression was confirmed in HEK293 cells 48 hours post-infection using Western blot analysis (Figure 1E).

### Systemic and mucosal immunogenicity studies in vaccinated mice

#### Humoral immune responses

H5CC vaccine immunogenicity was evaluated in female BALB/c mice using a heterologous prime-boost strategy. Mice (*n* = 10/group) were immunized with 2×10^10^ viral particles (vp) of Ad5_H5CC or Ad28_H5CC by combining intramuscular (IM) and intranasal (IN) coadministration. DPBS was used as a sham vaccine to serve as a negative control. Four weeks later, half of the mice (*n* = 5/group) were humanely sacrificed to assess prime immune responses. The remaining mice were boosted with an equivalent dose of Ad6_H5CC or Ad48_H5CC and sacrificed two weeks post boost to assess immune correlates (Figure 2A). Serum IgG responses were assessed via enzyme-linked immunosorbent assay (ELISA) against a panel of recombinant H5-HA proteins spanning from 1996 to 2024. Prime vaccination induced significant IgG responses in both vaccine groups compared to the DPBS control group (*p* < 0.0001). Ad5_H5CC consistently elicited higher serum IgG levels than the Ad28_H5CC group except against the closely related Vietnam/1194/04 strain (98.4% identity). Serotype-switched boosting further increased IgG titers across all six tested antigens, with both vaccine regimens remaining significantly higher than the DPBS control group (*p* < 0.0001) (Figure 2B).

**Figure 2:**
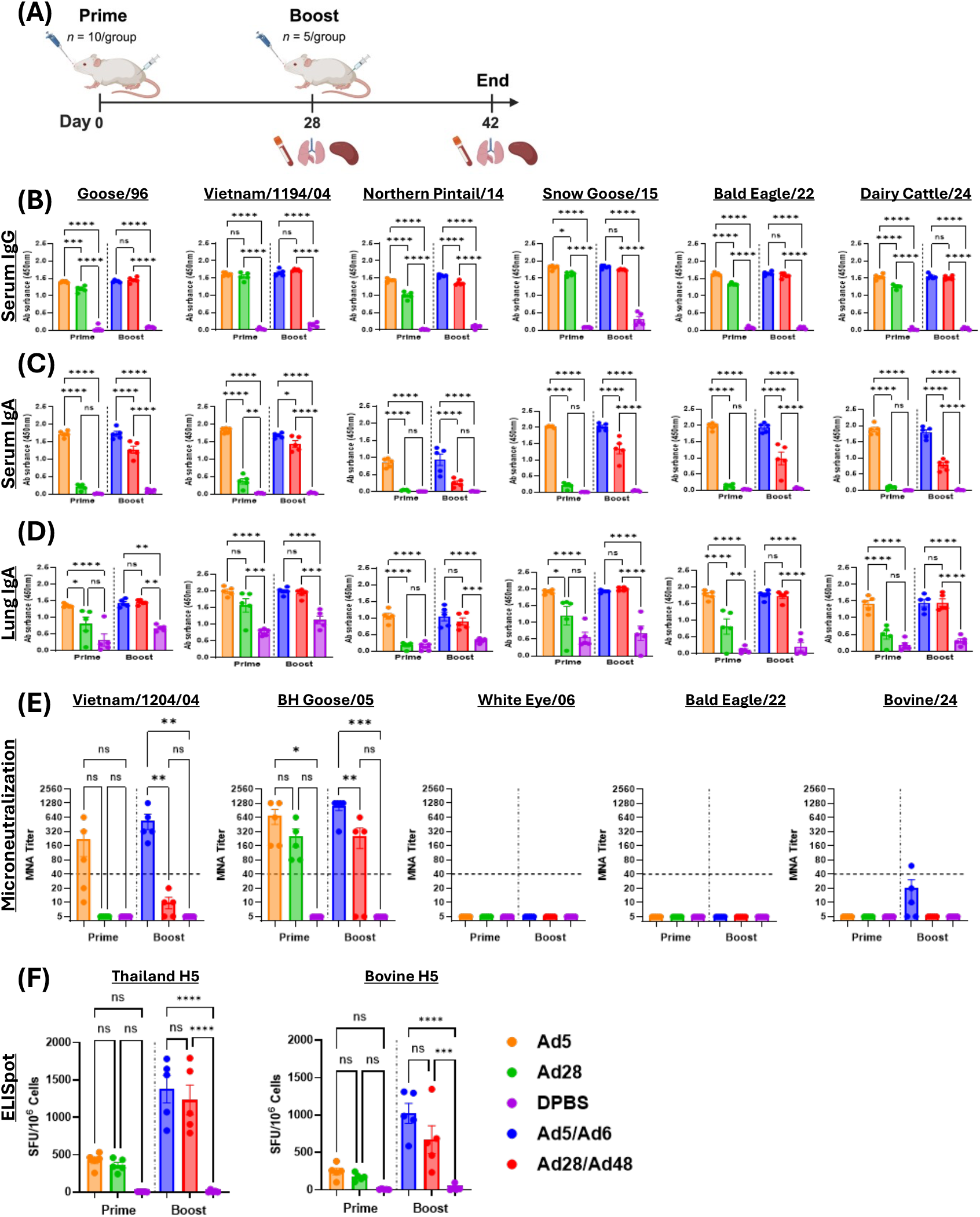
Mouse Humoral and Cellular Immune Correlates. (A); Schematic representation of mouse immunization timeline and tissue collection including serum, lung homogenates, and splenocytes. (B); Serum IgG ELISA. (C); Serum IgA ELISA. (D); Lung IgA ELISA. (E); Serum microneutralization assay (MNA). (F); Antigen specific T cell responses in splenocytes against H5-HA peptide libraries. Horizontal dashed line indicates 1:40 protective threshold. Statistical significance determined by one-way ANOVA with an alpha of 5.0%. \**p*<0.05, \*\**p*<0.01, \*\*\**p*<0.001, \*\*\*\**p*<0.0001; ns, not significant.

Circulating serum IgA analysis revealed the Ad5_H5CC vaccine consistently outperformed the Ad28_H5CC vaccine, with significantly higher antibody titers against all antigens compared to both the DPBS control group and the Ad28_H5CC vaccinated mice (*p* < 0.0001). In contrast, the Ad28_H5CC vaccine elicited a significant IgA response only against Vietnam/1194/04 (*p* < 0.01). Boost vaccination enhanced serum IgA titers, with the Ad5/Ad6_H5CC consistently showing elevated titers, while the Ad28/Ad48_H5CC induced significant increases against every antigen except Northern Pintail/14 (*p* < 0.0001) (Figure 2C). Mucosal IgA in lung supernatants mirrored systemic responses with the Ad5_H5CC vaccine inducing strong titers to five of the six tested antigens compared to the Ad28_H5CC vaccine. Ad5_H5CC also produced high mucosal IgA levels for all six HA antigens compared to the DPBS control group. Boost vaccination with Ad6_H5CC or Ad48_H5CC resulted in steadily high mucosal IgA in both vaccinated groups, with significant responses to all antigens compared to the DPBS control group (*p* < 0.01 or better). No significant differences were observed between the Ad5/Ad6 and the Ad28/Ad48 groups for any of the H5 antigens tested (Figure 2D).

Protective antibody functions were assessed through hemagglutination inhibition (HI) and microneutralization assays (MNA) against five H5 IAVs spanning 2004-2024. No sera from any of the vaccine groups produced detectable HI activity (Figure S2). In contrast, antibody neutralization improved when assessing sera using MNA. Notably, the Ad5_H5CC and Ad5/Ad6_H5CC groups exceeded the protective 1:40 threshold against Vietnam/1203/04 and BH Goose/05 (98.2% and 97.5% identity, respectively). The Ad5/Ad6_H5CC group also produced low, non-significant titers against Bovine/24, with a mean titer of 1:20. Ad28_H5CC prime vaccine reached protective titers against BH Goose/05 (1:256), while Ad28/Ad48_H5CC induced low neutralization against Vietnam/1203/04 (1:10) and robust protective titers against BH Goose/05 (1:258). The DPBS control group showed no neutralizing activity. None of the tested H5CC vaccines produced measurable MNA titers against the White Eye/06 or Bald Eagle/22 strains (Figure 2E).

#### Cell-mediated immune responses

To evaluate cell-mediated immunity induced by the Ad_H5CC vaccines, splenocytes were harvested from immunized mice following the timeline in Figure 2A. Cells were stimulated with two H5-HA peptide libraries (17-mers overlapping by 11 amino acids) derived from Thailand/04 (98.2% identity) and Bovine/24 (91.7% identity) (Figure S1). Antigen-specific T cell responses were quantified by interferon-gamma (IFN-γ) enzyme-linked immunospot (ELISpot) assay and expressed as spot forming units (SFU) per million cells. Prime-only vaccination elicited moderate responses: 426 and 246 SFU/10^6^ cells in the Ad5_H5CC group, and 360 and 171 SFU/10^6^ cells in the Ad28_H5CC group against Thailand/04 and Bovine/24, respectively. Heterologous boosting with the Ad6_H5CC or Ad48_H5CC increased spot counts ∼4-fold across both groups.

Ad5/Ad6_H5CC produced 1378 SFU/10^6^ and 1022 SFU/10^6^ cells against the Thailand/04 and Bovine/24 libraries, respectively, which were significantly higher than the DPBS control group (*p* < 0.0001). The species D Ad28/Ad48_H5CC group generated lower spot counts than the species C vaccines, (1232 SFU/10^6^ and 671 SFU/10^6^ cells), but still significantly exceeded the DPBS control group (*p* < 0.0001 and *p* < 0.001) (Figure 2F).

### Protection from lethal IAV H5N1 challenge in vaccinated mice

To assess vaccine-mediated protective efficacy, groups of BALB/c mice (*n* = 5) were immunized with Ad5/Ad6_H5CC or Ad28/Ad48_H5CC (2×10^10^ vp/animal, IM+IN) using a prime-boost regimen (Figure 3A). Two weeks post-boost, mice were challenged with lethal H5N1 reverse genetics (rg)-A/Vietnam/1203/2004 (Vietnam/1203/04) or rg-A/Bovine/Ohio/B24-OSU-439/2024 (Bovine/24), representing distinct antigenic diversity. Morbidity was monitored by daily weight measurements for 14 days post infection. Vaccinated mice showed minimal or no weight loss regardless of challenge strain (Figures 3B, 3C, and S3). Ad28/Ad48_H5CC vaccinated mice lost ∼2% body weight after Vietnam/1203/04 challenge, while Ad5/Ad6_H5CC maintained baseline weight (Figure 3B). Following Bovine/24 challenge, both vaccine groups maintained or exceeded baseline weight (100-105%) throughout the study (Figure 3C). In contrast, unvaccinated DPBS controls exhibited progressive weight loss beginning on day 2, reaching 25% by days 5-6 and requiring euthanasia. All vaccinated groups achieved 100% survival against both challenge viruses, whereas all control mice succumbed to infection (Figures 3D and 3E).

**Figure 3:**
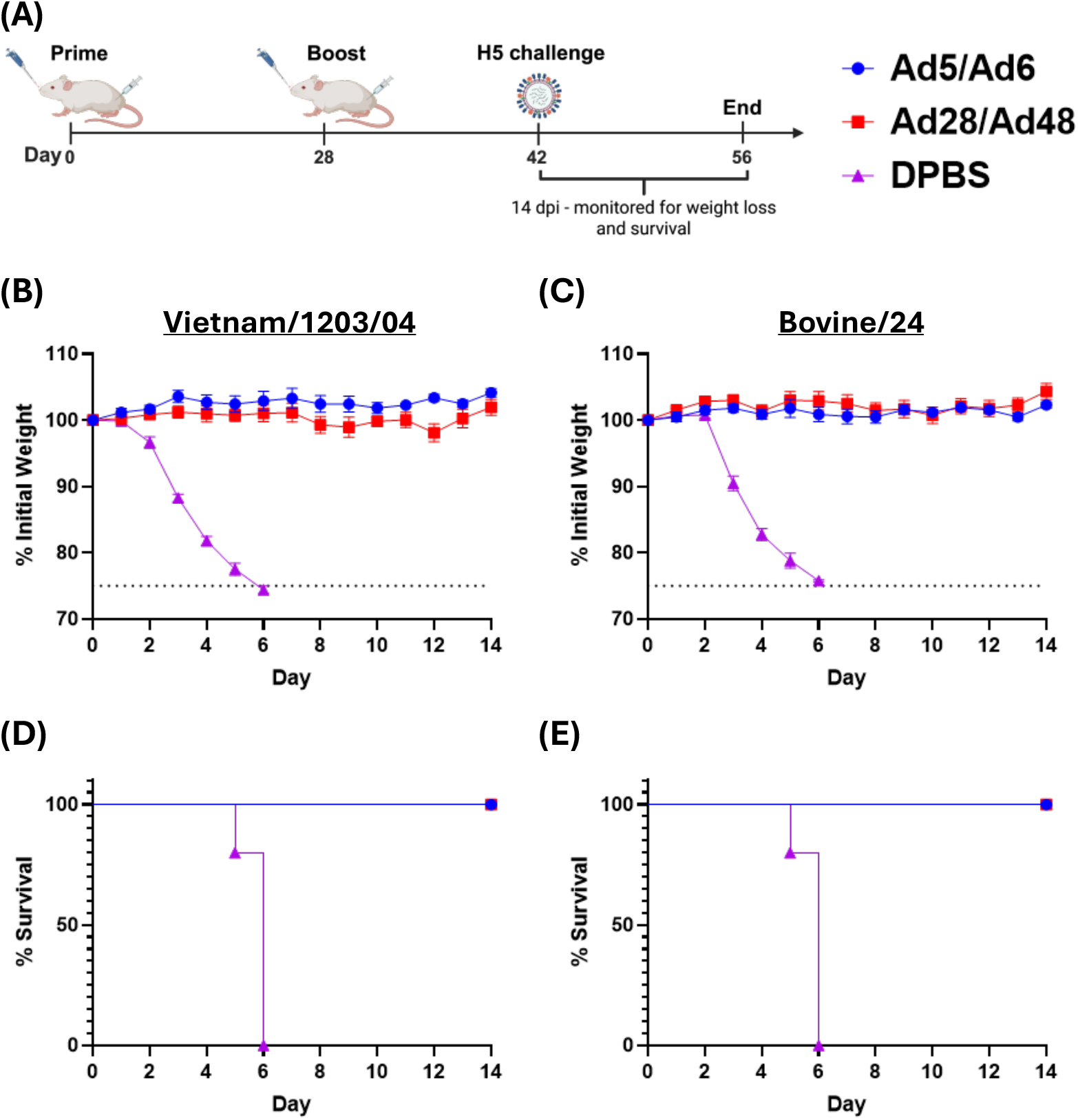
Protection against lethal IAV H5N1 challenge. (A); Schematic representation of vaccine and challenge timeline. Female BALB/c mice (*n* = 5/group) were vaccinated IM and IN with Ad-vectored H5CC on days 0 and 28. Infection with 100MLD_50_ of the indicated viruses occurred on day 42. (B,C); Mice were monitored for weight-loss and (D,E); survival for 14 days post-infection until the end of the study on day 56. Horizontal dashed line indicates 25% weight loss cutoff used for humane euthanasia.

### H5CC vaccine immunogenicity assessment in newborn calves

#### Humoral immune responses

Following confirmation of protective efficacy in mice, the immunogenicity of Ad-vectored H5CC vaccines was evaluated in Holstein dairy calves (*n* = 4/group) using a prime-boost regimen. Calves received an initial dose of either Ad5_H5CC or Ad28_H5CC (2×10^11^ vp/animal), via IM and IN coadministration. A mucosal atomization device was used for the IN vaccine delivery to ensure rapid adsorption of the vaccine across the mucosal epithelium. Blood and nasal swabs were collected before vaccination (day 0) to establish baseline immune responses, followed by weekly collections on days 7, 14, 28, 35, and 42. On day 28, calves were boosted with either Ad6_H5CC or Ad48_H5CC (2×10^11^ vp/animal) via IM and IN coadministration, and were euthanized two weeks later, on day 42 (Figure 4A).

**Figure 4:**
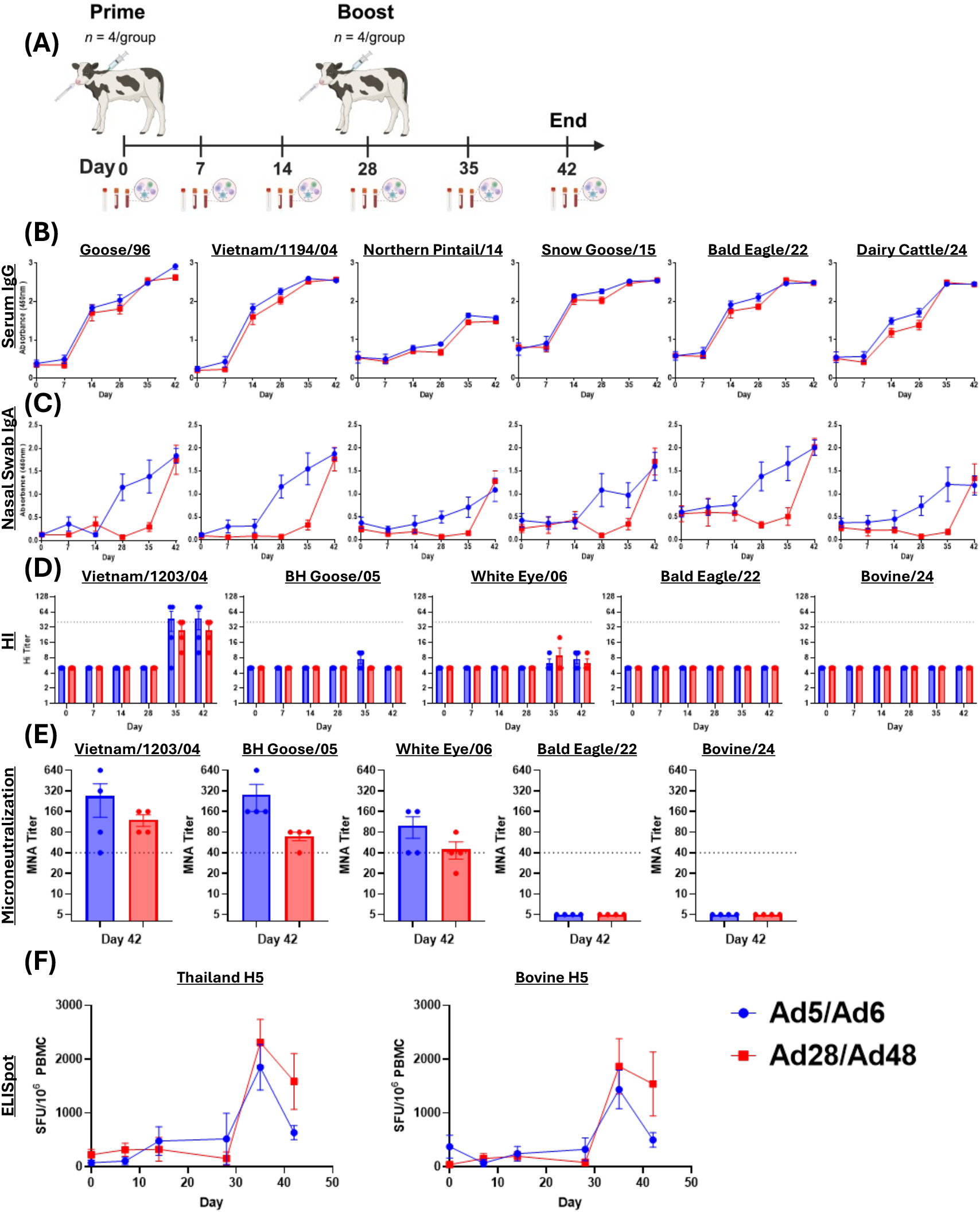
Immunized Calf Humoral and Cellular Immune Correlates. (A); Schematic representation of calf vaccine timeline and tissue collection including nasal swabs, serum, and PBMCs. (B); Serum IgG. (C); Nasal swab IgA. (D); Serum hemagglutination inhibition (HI) assay. (E); Serum microneutralization (MNA) assay. (F); Antigen specific PBMC T cell responses against H5-HA peptide libraries. Horizontal dashed line indicates 1:40 protective threshold.

Serum IgG responses were assessed by ELISA, against six H5-HA antigens, spanning from Goose/96 to the recent Dairy Cattle/24 strain. After prime vaccination, IgG levels were noticeably higher at day 14 for five of the six antigens (Goose/96, Vietnam/1194/04, Snow Goose/15, Bald Eagle/22, and Dairy Cattle/24) compared to the baseline. Boost vaccination further increased IgG levels, with both Ad6_H5CC and Ad48_H5CC groups reaching peak antibody titers against all antigens on day 42 (Figure 4B).

Mucosal IgA levels from nasal swabs remained near baseline through day 14 in the Ad5_H5CC group, then increased after the boost vaccination on day 28. By day 42, Ad6_H5CC boost-induced mucosal IgA levels were notably elevated against three antigens (Goose/96, Vietnam/1194/04, and Bald Eagle/22). The Ad28_H5CC group showed minimal mucosal IgA induction prior to boosting with Ad48_H5CC, after which a slight increase in IgA was observed on day 35 for Goose/96 and Vietnam/1194/04 antigens, and notable increases across all six antigens by day 42 (Figure 4C).

Functional antibody responses were evaluated via HI assay for five H5 influenza strains, including Bovine/24. Despite high ELISA titers on day 28, neither vaccine group showed measurable HI responses after the prime vaccination. The H5CC boost vaccines induced weak, but detectable HI titers against older strains (BH Goose/05 and White Eye/06) on days 35 and/or 42, but these were below the protective 1:40 threshold. HI titers against the more closely related Vietnam/1203/04 exceeded the 1:40 protective threshold on days 35 and 42 in the Ad5/Ad6_H5CC group (1:46 and 1:48, respectively). The Ad28/Ad48_H5CC group produced titers slightly below the protective threshold on both days 35 and 42 (1:28). No HI titers were observed for either group against the Bald Eagle/22 or Bovine/24 strains (Figure 4D).

Serum collected on day 42, demonstrating the highest total antibody levels measured by ELISA, was also tested using the more sensitive microneutralization assay (MNA). Both vaccine groups exceeded the 1:40 protective threshold for the Vietnam/1203/04, BH Goose/05, and White Eye/06 strains. The Ad5/Ad6_H5CC group had mean titers of 1:270, 1:280, and 1:100, while the Ad28/Ad48_H5CC group showed mean titers of 1:120, 1:70, and 1:45. Neither vaccine group achieved measurable neutralization titers against the Bald Eagle/22 or Bovine/24 strains (Figure 4E).

#### Cell based immunity

Cross-reactive T cell responses were assessed by IFN-γ ELISpot using PBMCs stimulated with two H5-HA peptide libraries: Thailand/04 (98.2% identity) and Bovine/24 (91.7% identity), each consisting of 17-mer peptides overlapping by 11 amino acids. Prime vaccination did not induce significant antigen-specific T cell responses, with spot counts remaining near baseline levels through day 28 in both vaccine groups. However, Ad6_H5CC or Ad48_H5CC boost-vaccination resulted in notable increases in H5-specific T cell responses to both libraries, peaking one-week post-boost at day 35. The Ad28/Ad48_H5CC group induced the highest responses, with mean spot counts of 2314 and 1870 SFU/10^6^ cells at day 35 for the Thailand/04 and Bovine/24 libraries, respectively. In comparison, the Ad5/Ad6_H5CC group had lower responses on day 35 (1851 and 1440 SFU/10^6^ cells) but still exhibited higher T cell responses than baseline. T cell responses declined in both vaccine groups two-weeks post-boost (day 42) but remained elevated compared to day 0 (Figure 4F).

## Discussion

The emergence of IAV in cattle is a recent phenomenon, as they were not considered a host species prior to the 2024 HPAI H5N1 outbreak (26). The mechanisms of interspecies transmission from wild birds to cattle, and subsequent spread both within cattle populations and to humans, remain under investigation. However, nasal shedding does not appear to play a major role (10, 14). Based on current influenza knowledge and understanding, limiting IAV transmission will likely require vaccines capable of eliciting broad, durable immune responses. Currently, no licensed influenza vaccines exist for cattle. Adenoviral (Ad)-vectored platforms represent a promising approach, as they are well characterized as influenza vaccines (27, 28). In cattle, bovine Ad3 and human Ad5 vectors have been evaluated against non-influenza pathogens, eliciting robust immune responses (29, 30). While species C adenoviruses have demonstrated immunogenicity in cattle, species D adenoviruses have not been evaluated prior to this study. Their ability to bind bovine receptors such as bovCD46 and mediate efficient cell entry is currently unknown, possibly explaining their overall reduced immunogenicity relative to species C vectors.

In addition to systemic immunity, mucosal immunity is critical for preventing respiratory epithelium infection and for limiting viral transmission (31-33). To induce both responses, we employed a dual-route, synchronous vaccination strategy combining intramuscular and intranasal administration. A prime–boost regimen was employed with serotype-switched species C (Ad5/Ad6) or species D (Ad28/Ad48) vectors, as heterologous Ad serotype switching has previously been shown to enhance immune responses compared to homologous boosting (20, 23, 34-38). In both mice and cattle, the H5CC vaccine induced strong, cross-reactive systemic and mucosal antibody responses against historical clade 0 and clade 1 strains, as well as the more recently emerged avian and bovine clade 2.3.4.4b strains. Although hemagglutinin inhibition (HI) and microneutralization (MNA) titers against clade 2.3.4.4b viruses were modest, vaccinated mice were completely protected against weight loss and death after high-dose lethal challenge. These findings suggest that breadth and magnitude of antibody responses elicited in cattle could translate into meaningful protection. Protection from challenge may involve Fc-mediated antibody effector functions such as antibody-dependent cellular cytotoxicity (ADCC) or T cell-mediated immunity, consistent with prior influenza vaccine studies (39-41).

T cell responses are also critical for protection and viral clearance. Adenoviral vectors are advantageous because they efficiently deliver transgenes without adjuvant, inducing long-lived T cell immunity, especially when using rare serotypes such as species D adenoviruses, which have low pre-existing seroprevalence. In our study, both species C (Ad5/Ad6) and species D (Ad28/Ad48) vectors induced strong IFN-γ secreting T cell responses under a prime–boost regimen, with Ad28/Ad48 generating the most robust responses in calves. The species D vectors were E3-deleted and replication competent, possibly explaining the durability of immunity and higher T cell responses compared to species C (replication defective) vectors in cattle. Importantly, vaccine-induced T cells responded to peptide libraries derived from both an older Thailand 2004 strain and the currently circulating clade 2.3.4.4b bovine 2024 strain. This suggests that these vaccines could aid in controlling ongoing HPAI H5 outbreaks.

This study has some limitations. First, the vaccine design excluded the multibasic cleavage site, and challenge experiments were performed with genetically modified, reassorted highly-pathogenic avian influenza (HPAI) viruses in order to create low-pathogenic avian influenza (LPAI) viruses suitable for BSL-2 conditions. It is therefore unknown whether the vaccine would provide equivalent protection against systemic viremia and severe disease caused by HPAI challenge. However, the lack of detectable disease in the mouse model indicates that the infection is rapidly prevented. If this is the case, H5CC vaccine induced immunity could halt viral dissemination that is conferred by increased tropism inherent to HPAI viruses. Second, only male calves were available for testing, leaving potential sex-based differences in vaccine immunogenicity untested. Third, we did not perform cattle challenge studies with H5Nx viruses. Thus, the extent to which our vaccine construct would prevent infection, disease, or shedding in cattle infection remains unknown. Therefore, future work should assess vaccine efficacy and sex-based immunogenicity in the target species by measuring viral RNA titers and histopathology in respiratory tissues, as well as in milk, to ensure safety and efficacy.

This study evaluated a H5 consensus vaccine, derived from sequences over 20 years old, to assess its cross-reactivity against the contemporary 2.3.4.4b HPAI H5Nx strains and evaluate its immunogenicity in dairy cattle. Our data demonstrate that a vaccine originally designed for human H5 strains has strong potential against newly circulating bovine strains. The H5CC vaccine induced potent humoral and cellular immunity, and the dual-route administration strategy successfully elicited both systemic and mucosal responses. Given the absence of licensed influenza vaccines for cattle, a broad-spectrum adenoviral-vectored vaccine platform is a promising strategy to decrease economic losses in the livestock sector and reduce the risk of H5 influenza transmission within cattle populations and from cattle to humans.

## Materials and Methods

### Ethics statement

All biological experiments conducted in this study were approved by the University of Nebraska – Lincoln Institutional Biosafety Committee (IBC) under protocol number 619. All experiments involving mice and calves were further approved by the Institutional Animal Care and Use Committee (IACUC) under protocols 2662, and 2717 respectively. Experiments were carried out under BSL-2+ conditions according to the NIH guide for the Care and Use of Laboratory Animals. All animals used in this study were allowed to acclimate for one week prior to immunization and were provided access to food and water *ad libitum*. Female BALB/c mice were purchased from Jackson Laboratories and housed in groups of 5 mice per cage in a temperature-controlled room with a 14-hour light, 10-hour dark cycle. Eight one-week-old, male Holstein dairy calves were split into two equal groups of 4 per room. The calves were housed on a 14-hour light, 10-hour dark cycle. Calves were bottle fed with Purina Land O Lakes Herd Maker Protein Blend twice per day until 6 weeks of age, at which point milk feeding was decreased to once per day. Pellet food was introduced into their diet beginning at 4 weeks of age. Calves were fed appropriate rations for their species and age.

### Cells and viruses

Human Embryonic Kidney 293 (HEK293) cells and Madine-Darby Canine Kidney (MDCK) cells were obtained from the American Type Culture Collection (ATCC). Cells were propagated in Dulbecco’s Modified Eagle Medium (DMEM) supplemented with 10% fetal bovine serum (FBS) and 1% penicillin-streptomycin (P/S) and cultured in a humidified incubator at 37 °C with 5% CO_2_. A BSL-2 compliant reverse genetic (rg) system was used to produce the following H5N1 Influenza A virus (IAV) strains for use in this study: rg-A/Vietnam/1203/2004 (Vietnam/1203/04), rg-A/BH Goose/Qinghai/A/2005 (BH Goose/05), rg-A/Japanese White Eye/Hong Kong/1038/2006 (White Eye/06), rg-A/Bald Eagle/Florida/W22-134-OP/2022 (Bald Eagle/22), and rg-A/bovine/Ohio/B24OSU-439/2024 (Bovine/24). Six (PB1, PB2, PA, NP, M, and NS) IAV gene segments from the PR/8/34 H1N1 laboratory strain were cloned individually into the pHW2000 vector (9, 42). Separately, the neuraminidase (N) gene and hemagglutinin (H) gene without the highly pathogenic multibasic cleavage site from each strain were synthesized and cloned into the same pHW2000 vector. All 8 plasmids (250 ng each) were transfected using Polyfect™ into a 6-well plate seeded with a 50:50 mixture of HEK293 and MDCK cells. Viral supernatants were collected 3 days post-transfection and used to infect 11-day old premium plus specific pathogen free embryonated chicken eggs (AVSbio). Three days later, allantoic fluid was harvested from the eggs and tested for hemagglutination units (HAU) titer (described below). Allantoic fluid was harvested 3 days later, tested for hemagglutination units, pooled by strain, split into single-use aliquots, and stored at -80 °C.

### Vaccine design

Twenty-one distinct H5-HA sequences collected from 1997 to 2005 were used to create the H5 centralized consensus (H5CC) vaccine. The full length H5 gene sequences were downloaded from the Influenza Research Database and aligned using ClustalX v2.1, generating a 568 amino acid consensus sequence as previously published (23). The H5CC sequence was codon-optimized for humans and synthesized by GenScript. The synthesized gene was cloned into the pFastBac-1 plasmid to facilitate recombinant vaccine production as described below.

### Adenovirus production

Four recombinant human adenoviruses representing species C (type 5 and type 6) and species D (type 28 and type 48) were used in this study. These serotypes encompass high-seroprevalent (species C) and low-seroprevalent (species D) subtypes. The H5CC gene derived from the pFastBac-1 vector was cloned into the Ad5 pShuttle-CMV vector (or analogous vectors for Ad6, Ad28, and Ad48) through digestion with *KpnI* and *HindIII* restriction enzymes. The resulting shuttle plasmid was mixed with pAdEasy (Agilent Technologies) and co-transformed into BJ5183 *E. coli* cells to facilitate homologous recombination of the H5CC gene into the adenoviral genome. The recombinant adenoviral genomes were linearized via *AscSI* restriction digestion and transfected into HEK293 cells in a 6-well plate. Viral constructs were amplified through sequential passages in HEK293 cells and propagated in a Corning 10-cell stack to achieve a high viral titer. After 72 hours, infected cells were harvested, pelleted by centrifugation, and the recombinant adenoviruses were purified through two rounds of CsCl ultracentrifugation as previously described (43, 44). Viral stocks were desalted using a Bio-Rad Econo-Pac 10DG column, titered by measuring the OD_260_, split into aliquots, and frozen at -80 °C for long-term storage.

### Adenoviral Vector Sequencing

Recombinant adenoviral DNA was extracted according to the manufacturer’s instructions using a Purelink viral RNA/DNA Mini kit (Invitrogen). Extracted viral DNA (vDNA) was eluted in water and quantified using a NanoDrop spectrophotometer. Oxford Nanopore sequencing was conducted following the Amplicon Sequencing from DNA protocol. Briefly, 200 ng vDNA was diluted to a final volume of 10 µL in water, mixed with 1.5 µL of rapid barcode, and transferred to a thermocycler set for 2 min at 30 °C, followed by 2 min at 80 °C. Barcoded samples were pooled, mixed with an equal volume of AMPure XP magnetic beads for 10 min on a rotator at room temperature, and pelleted by centrifugation. The DNA library was washed three times with 80% ethanol on a magnetic rack and eluted in 15 µL elution buffer for 10 min. Five microliters of pooled, barcoded DNA was mixed with rapid adapter (RA), RA buffer, sequencing buffer, and library beads according to the protocol, loaded onto a Flongle, and sequenced with a MinION Mk1B. High quality sequencing reads were assembled using the Amplicon workflow in Epi2Me version 5.2.4.

### Western Blotting

To confirm Ad-vectored H5CC protein expression, Western blotting was performed as previously described (44). In short, HEK293 cells were seeded in a 12-well plate with cDMEM and cultured overnight at 37 °C with 5% CO_2_. The following day, cells were infected with Ad-vectors at an MOI of 100 for 48 hours. Cells were harvested in Laemmli buffer (BioRad) with 10% β-mercaptoethanol, lysed at 95 °C for 10 min, and passed through a QIAShredder (QIAgen). Lysates were loaded onto a 12% SDS-PAGE gel and resolved at 150 V for 2 hours. Proteins were transferred onto a nitrocellulose membrane for 1 hour at 60 V, followed by blocking for 1 hour in Tris-Buffered Saline + 0.1% Tween-20 (TBST) with 5% skim milk. To detect the recombinant HA protein, the membrane was incubated overnight at 4 °C with polyclonal goat serum raised against A/tern/South Africa/1961 (BEI, NR-3156) diluted 1:1000 in TBST + 2% skim milk. After washing with TBST, the membrane was probed for 1 hour with a donkey anti-goat IgG-HRP secondary antibody (R&D systems HAF109) diluted 1:5000 in TBST + 2% skim milk. Following additional TBST washes, the membrane was developed for 10 min using SuperSignal West Pico Chemiluminescent substrate (ThermoFisher) and imaged on a ChemiDoc MP system (BioRad). The membrane was stripped for 30 min in a pH 2.2 solution containing 15% (w/v) glycine, 1% (w/v) SDS, 1% (v/v) Tween 20. After stripping, the membrane was washed thoroughly with TBST and re-probed overnight using a mouse anti-GAPDH-HRP conjugated antibody (Santa Cruz, sc-47724) diluted 1:1000 in TBST + 2% skim milk. The membrane was developed as described above.

### Mouse Immunizations

Mice were immunized through prime-boost vaccination with 2×10^10^ viral particles (vp) coadministered through intramuscular (IM) and intranasal (IN) delivery. Prior to vaccination, mice were anesthetized via ketamine/xylazine intraperitoneal injection. For IM-immunization, a total volume of 50 µL was injected, with 25 µL injected into each hind leg muscle. For IN-immunization, the total volume was 20 µL, with 10 µL administered by pipette into each nostril. Sterile DPBS was used as a mock vaccine. Boost vaccines were administered on day 28 via the same procedure. On days 28 or 42 post-vaccination, selected groups of mice were humanely euthanized, and their blood, lungs, and spleens were harvested to assess immune responses. The remaining groups were anesthetized with ketamine/xylazine and challenged intranasally with 100 times the median lethal dose (100MLD_50_) of the indicated H5 IAV strains to assess protection against lethal challenge. Challenged mice were monitored daily for weight loss for 14 days. Mice were humanely euthanized if they lost ≥25% body weight.

### Calf Immunizations

Calves were immunized through prime-boost vaccination with 2×10^11^ vp/animal (IM+IN coadministration) on days 0 and 28. IM-vaccination was administered via syringe to the neck muscle in a total volume of 1 mL. IN-vaccination was administered using a MAD Nasal™ Intranasal Mucosal Atomization Device (Teleflex) attached to a syringe, with a dose of 500 µL sprayed into each nostril. Blood samples were collected on days 0, 7, 14, 28, 35, and 42 in BD Vacutainer Serum Separator tubes and EDTA Vacutainer tubes (Benton Dickenson) for serum and peripheral blood mononuclear cells (PBMC) isolation. Nasal swabs were collected in UniTranz-RT Universal Transport Medium tubes (Puritan) to assess mucosal IgA levels. Calves were humanely euthanized on day 42 under the care of the UNL veterinary staff with a mixture of ketamine/xylazine/butorphanol for sedation followed by a lethal dose of pentobarbital.

### Splenocyte and PBMC Isolation

Mouse spleens were surgically excised, transferred to 15 mL tubes containing RPMI-1640 (Gibco) media supplemented with 5% FBS and 1% P/S, hereafter referred to as complete RPMI, and stored on ice. Spleens were mechanically disrupted by repeated passage through 40 µm cell strainers, followed by red blood cell lysis with ACK lysis buffer. Splenocytes were then washed with complete RPMI media and immediately counted for ELISpot assays or pelleted and resuspended in 90% FBS and 10% DMSO freezing media for long-term cryopreservation. As previously mentioned, bovine PBMCs were isolated by first collecting whole blood in BD Vacutainer tubes containing EDTA. Whole blood was diluted 1:1 in sterile DPBS, and gently layered over Lymphocyte Separation Media (Corning), followed by centrifugation at 1200 *g* for 45 min. The lymphocyte layer was carefully harvested from the mixture using a serological pipette, treated with ACK lysis buffer and resuspended in complete RPMI or freeze media as outlined above.

### T cell Responses

Enzyme-linked immunospot (ELISpot) assays were conducted on mouse splenocytes or bovine PBMCs to assess vaccine-induced, antigen-specific interferon gamma (IFN-γ) T cell responses. 96-well PVDF-backed plates were coated overnight at 4 °C with 50 µL/well of 5 µg/mL IFN-γ monoclonal antibody (mouse: MabTech AN18; bovine: MabTech MT17.1). The following day, plates were washed with DPBS and blocked with 250 µL complete RPMI at 37 °C. Fresh splenocytes/PBMCs were resuspended directly in complete RPMI. Frozen samples were thawed at 37 °C for 2 min, washed, and resuspended in complete RPMI. Live cells were counted on a Bio-Rad TC20 counter with Trypan blue and adjusted to 2.5×10^4^ cells/µL. One hundred microliters of cell suspension were seeded in duplicate wells. Cells were stimulated with overlapping H5-HA peptide libraries (17-mer peptides, 11 amino acid overlap): A/Thailand/4(SP-528)/2004 (BEI Resources, NR-2604), or Bovine/24 (GenScript). A 50 µL peptide pool containing 250 ng of each peptide was then added to each well. Concanavalin-A (5 µg/mL; Sigma) and complete RPMI served as positive and negative controls, respectively. Plates were incubated at 37 °C with 5% CO_2_ for 16-18 hours. Wells were then washed 6 times with DPBS + 0.1% Tween-20 (DPBS-T) and incubated for 1 hour at room temperature with 50 µL of a DPBS + 1% FBS containing 1 µg/mL of biotinylated monoclonal detecting antibody (mouse: MabTech R4-6A2; bovine: MabTech MT307). After additional DPBS-T washes, plates were developed using BCIP/NBT (Plus) alkaline phosphatase substrate (Thermo-Fisher). Development was stopped by washing several times in ddH_2_O. Plates were air-dried overnight before counting on an automated ELISpot reader (AID iSpot Reader Spectrum, AID GmbH). Results were normalized with negative controls and expressed as spot-forming units (SFU) per 10^6^ splenocytes or PBMCs.

### HI Assay

One volume of mouse or calf serum was incubated overnight at 37 °C with three volumes of receptor destroying enzyme (Hardy Diagnostics #370013). The following day treated serum was heat-inactivated at 56 °C for 30 min and adjusted to a final ratio of 1:10 with six volumes of DPBS. Fifty microliters of diluted serum were transferred to the top row of 96-well V-bottom plates, and two-fold serial dilutions (25 µL/well) were performed. Next, 8 HAU of the indicated H5 viruses, diluted to 25 µL in DPBS were added to each well. Serum and virus mixtures were incubated at room temperature for 1 hour. Afterwards, 50 µL of rooster red blood cells (RBCs; Lampire Biologicals) at a concentration of 4×10^7^ cells/mL were added to each well. Plates were incubated for 45 min at room temperature. Hemagglutination inhibition (HI) was assessed by tilting the plates at a 45° angle to observe teardrop formation. The highest dilution of serum that produced teardrops was recorded as the HI titer; if no teardrops were observed, an HI titer of 5 was recorded. Negative controls (virus only, no serum) and positive controls (no virus, no serum) were included in each assay, and virus back titration was performed to confirm assay validity.

### Microneutralizations

Mouse or calf serum was heat-inactivated at 56 °C for 30 min, then 50 µL of serum was transferred to the top row of 96-well U-bottom plates. Two-fold serial dilutions (25 µL/well) were prepared in DPBS, followed by the addition of 100 tissue culture infective dose 50% (TCID_50_) of the specific H5 viruses in a volume of 25 µL/well. Plates were incubated for 1 hour at 37 °C with 5% CO_2_. Subsequently, 200 µL of MDCK cells (2×10^5^ cells/mL) were added to each well and incubated overnight at 37 °C and 5% CO_2_. The following day, plates were washed with DPBS and 200 µL/well of DMEM supplemented with 2 µg/mL TPCK-treated trypsin was added to each well. Plates were returned to the incubator for 3 days. Next, 50 µL of 4×10^7^ rooster RBCs/mL were added to each well, and plates were incubated at room temperature for 1 hour. Hemagglutination was assessed by tilting the at a 45° angle as described above. Positive and negative controls, as well as back-titrations, were included in each assay to confirm validity.

### ELISA

A panel of six recombinant H5 antigens spanning from 1996 to 2024 and representing clades 0, 1, 2.3.4.4c, and 2.3.4.4b (avian and dairy cattle) were utilized to assess antibody responses induced by the H5CC vaccines using enzyme-linked immunosorbent assay (ELISA). The H5-HA antigens utilized were: A/goose/Guangdong/1/1996 (Goose/96; Sino Biological, Cat# 40024-V08H1), A/Vietnam/1194/2004 (Vietnam/1194/04; Sino Biological, Cat# 11062-V08H1), A/northern pintail/Washington/40961/2014 (Northern Pintail/14; BEI Resources NR-50174), A/snow goose/Missouri/CC15-84A/2015 (Snow Goose/15; BEI Resources NR-50651), A/bald eagle/Florida/W22-134-OP/2022 (Bald Eagle/22; BEI Resources NR-59922), and A/dairy cattle/Texas/24-008749-001-original/2024 (Dairy Cattle/24; BEI Resources NR-59816). Microlon high-binding 96-well plates (Greiner 655081) were coated with 200 ng of antigen in a final volume of 50 µL bicarbonate/carbonate coating buffer and incubated overnight at 4 °C. The following day, plates were washed multiple times with DPBS-T and blocked for 2 hours with 250 µL of DPBS-T + 4% bovine serum albumin (BSA). After additional washes, 50 µL of lung homogenate or nasal swabs (diluted 1:4 with DPBS containing dithiothreitol), or serum (diluted 1:100 in DPBS-T + 2% BSA), was added to the wells and incubated for 2 hours at room temperature. After incubation, plates were washed again and 50 µL of an appropriate secondary antibody (sheep anti bovine IgG: Bethyl A10-118P; rabbit anti bovine IgA: Bethyl A10-108P; goat anti Mouse IgA: Southern Biotech 1040-05; and goat anti mouse IgG: Millipore Sigma AP308P) conjugated with HRP diluted at 1:5000 in DPBS-T + 2% BSA was added to each well for 1 hour. After a final wash step 50 µL of 1-Step Ultra TMB-Substrate was added to each well until color formation. The reaction was stopped with an equal volume of 2 M sulfuric acid. Absorbance was measured on a SpectraMax i3X plate reader at 450nm. Wells treated without sera, lung homogenates, or nasal swabs, were used as a negative control and this background absorbance was subtracted from sample values.

### Statistical Analysis

Statistical analysis was conducted using GraphPad Prism version 10.6. Data are expressed as mean with standard error (mean ± SEM). Murine HI, MNA, ELISA, and ELISpot data were analyzed by one-way analysis of variance (ANOVA) with Tukey’s multiple comparisons. Survival outcomes were analyzed using the Kaplan-Meier log-rank test. A *p*-value below 0.05 was considered statistically significant.

## Acknowledgments

The authors would like to thank the veterinarians and technicians at the University of Nebraska – Lincoln Life Science Annex for their support and care of the animals used in this study. The following reagents were obtained through BEI Resources, NIAID, NIH: H5 Hemagglutinin (HA) Protein from Influenza Virus, A/northern Pintail/WA/40964/2014 (H5N8), Recombinant from Baculovirus, NR-50174; H5 Hemagglutinin (HA) Protein from Influenza Virus, A/snow goose/Missouri/CC15-84A/2015 (H5N2), Recombinant from Baculovirus, NR-50651; H5 Hemagglutinin (HA) Protein from Influenza A Virus, A/bald eagle/Florida/W22-134-OP/2022 (H5N1), Recombinant from Baculovirus, NR-59922; H5 Hemagglutinin (HA) Protein from Influenza A Virus, A/dairy cattle/Texas/24-008749-001-original/2024 (H5N1), Recombinant from Baculovirus, NR-59816; Peptide Array, Influenza Virus A/Thailand/4(SP-528)/2004 (H5N1) Hemagglutinin Protein, NR-2604; and Polyclonal Anti-Influenza Virus H5 (Hav5) Hemagglutinin (HA), A/tern/South Africa/1961 (H5N3) (antiserum, Goat), NR-3156. We would also like to thank Dr. Richard Webby at St. Jude Children’s Research Hospital, Memphis, TN for providing the reverse genetics H5N1 viruses used in this study.

