## Supplemental Table 1 for "Dual-Route H5N1 Vaccination Induces Systemic and Mucosal Immunity in Murine and Bovine Models"

### Bovine H5CC Manuscript

Supplemental Figures (3 total)

| | H5CC | A/Viet_... | A/Viet_... | A/Viet_... | A/Viet_... | A/Viet_... | A/Viet_... | A/Viet_... | A/Thaila... | A/Thaila... | A/Thaila... | A/Thaila... | A/Thaila... | A/Thaila... | A/snow... | A/Prach... | A/north... | A/Japan... | A/Hong... | A/Hong... | A/Hong... | A/Hong... | A/Hong... | A/Hong... | A/Hatay... | A/Hano... | A/goos... | A/dairy ... | A/Camb... | A/Bovin... | A/bar h... | A/bald ... |
|---|---|---|---|---|---|---|---|---|---|---|---|---|---|---|---|---|---|---|---|---|---|---|---|---|---|---|---|---|---|---|---|---|
| H5CC | | 97.0% | 98.1% | 97.9% | 97.9% | 98.1% | 98.2% | 98.4% | 96.1% | 97.9% | 98.1% | 98.2% | 97.9% | 98.2% | 92.1% | 98.4% | 91.9% | 95.7% | 95.6% | 95.4% | 95.4% | 94.9% | 97.7% | 97.6% | 97.5% | 98.1% | 96.0% | 91.7% | 97.5% | 91.7% | 97.5% | 91.5% |
| A/Viet_Nam/JPHN30321/... | 97.0% | | 98.6% | 98.4% | 97.7% | 98.2% | 98.4% | 98.6% | 95.6% | 98.4% | 98.2% | 98.4% | 98.1% | 98.4% | 91.9% | 98.6% | 91.7% | 95.2% | 96.0% | 95.4% | 95.4% | 95.1% | 97.4% | 97.3% | 97.7% | 97.9% | 95.6% | 91.4% | 98.2% | 91.4% | 95.8% | 91.2% |
| A/Viet_Nam/JP4207/2005 | 98.1% | 98.6% | | 99.5% | 98.8% | 99.3% | 99.5% | 99.6% | 96.7% | 99.1% | 99.3% | 99.5% | 99.1% | 99.5% | 92.1% | 99.6% | 91.9% | 95.8% | 95.8% | 96.0% | 96.0% | 95.1% | 98.1% | 98.0% | 98.8% | 98.9% | 96.1% | 91.5% | 99.1% | 91.5% | 95.8% | 91.4% |
| A/Viet_Nam/JP14/2005 | 97.9% | 98.4% | 99.5% | | 98.6% | 99.1% | 99.3% | 99.5% | 96.5% | 98.9% | 99.1% | 99.3% | 98.9% | 99.3% | 92.3% | 99.5% | 92.1% | 95.6% | 95.6% | 95.4% | 95.4% | 94.9% | 97.9% | 97.8% | 98.6% | 98.8% | 96.0% | 91.7% | 98.9% | 91.7% | 95.6% | 91.5% |
| A/Viet_Nam/HN30408/2... | 97.9% | 97.7% | 98.8% | 98.6% | | 98.8% | 98.9% | 99.1% | 96.1% | 98.6% | 98.8% | 98.9% | 98.6% | 98.9% | 91.9% | 99.1% | 91.7% | 95.4% | 95.4% | 95.2% | 95.2% | 94.7% | 97.5% | 97.5% | 98.2% | 99.1% | 95.8% | 91.4% | 98.2% | 91.4% | 95.8% | 91.2% |
| A/Viet_Nam/3046/2004 | 98.1% | 98.2% | 99.3% | 99.1% | 98.8% | | 99.5% | 99.6% | 96.8% | 99.1% | 99.3% | 99.5% | 99.1% | 99.5% | 92.4% | 99.6% | 92.2% | 95.8% | 95.8% | 95.6% | 95.6% | 95.1% | 98.1% | 98.0% | 98.8% | 98.9% | 96.1% | 91.5% | 98.9% | 91.5% | 95.8% | 91.3% |
| A/Viet_Nam/1203/2004 | 98.2% | 98.4% | 99.5% | 99.3% | 98.9% | 99.5% | | 99.8% | 97.0% | 99.3% | 99.5% | 99.6% | 99.3% | 99.6% | 92.2% | 99.8% | 92.0% | 95.9% | 95.9% | 95.8% | 95.8% | 95.2% | 98.2% | 98.2% | 98.9% | 99.1% | 96.3% | 91.7% | 99.1% | 91.7% | 95.9% | 91.5% |
| A/Viet_Nam/1194/2004 | 98.4% | 98.6% | 99.6% | 99.5% | 99.1% | 99.6% | 99.8% | | 97.0% | 99.5% | 99.6% | 99.8% | 99.5% | 99.8% | 92.4% | 100% | 92.3% | 96.1% | 96.1% | 96.0% | 96.0% | 95.4% | 98.4% | 98.4% | 99.1% | 99.3% | 96.5% | 91.9% | 99.1% | 91.9% | 96.1% | 91.7% |
| A/Thailand/LFPN_2004/2... | 96.1% | 95.6% | 96.7% | 96.5% | 96.1% | 96.8% | 97.0% | 97.0% | | 96.5% | 96.7% | 96.8% | 96.5% | 96.8% | 89.8% | 97.0% | 90.0% | 94.0% | 94.0% | 93.8% | 93.8% | 93.3% | 95.8% | 96.6% | 96.5% | 96.5% | 94.4% | 89.6% | 96.3% | 89.6% | 94.0% | 89.4% |
| A/Thailand/Chaiyaphum... | 97.9% | 98.4% | 99.1% | 98.9% | 98.6% | 99.1% | 99.3% | 99.5% | 96.5% | | 99.1% | 99.3% | 98.9% | 99.3% | 92.3% | 99.5% | 92.1% | 95.8% | 96.1% | 95.6% | 95.6% | 95.4% | 97.9% | 97.8% | 98.6% | 98.8% | 96.3% | 91.5% | 98.8% | 91.5% | 96.0% | 91.4% |
| A/Thailand/16/2004 | 98.1% | 98.2% | 99.3% | 99.1% | 98.8% | 99.3% | 99.5% | 99.6% | 96.7% | 99.1% | | 99.5% | 99.8% | 99.5% | 92.3% | 99.6% | 92.1% | 95.8% | 95.8% | 95.6% | 95.6% | 95.1% | 98.1% | 98.0% | 98.8% | 98.9% | 96.1% | 91.7% | 98.8% | 91.7% | 95.8% | 91.5% |
| A/Thailand/4(SP-528)/20... | 98.2% | 98.4% | 99.5% | 99.3% | 98.9% | 99.5% | 99.6% | 99.8% | 96.8% | 99.3% | 99.5% | | 99.3% | 99.6% | 92.3% | 99.8% | 92.1% | 95.9% | 96.0% | 95.8% | 95.8% | 95.3% | 98.2% | 98.2% | 98.9% | 99.1% | 96.3% | 91.7% | 98.9% | 91.7% | 96.0% | 91.6% |
| A/Thailand/2(SP-33)/2004 | 97.9% | 98.1% | 99.1% | 98.9% | 98.6% | 99.1% | 99.3% | 99.5% | 96.5% | 98.9% | 99.8% | 99.3% | | 99.3% | 92.1% | 99.5% | 91.9% | 95.6% | 95.6% | 95.4% | 95.4% | 94.9% | 97.9% | 97.8% | 98.6% | 98.8% | 96.0% | 91.5% | 98.6% | 91.5% | 95.6% | 91.4% |
| A/Thailand/1(KAN-1)/20... | 98.2% | 98.4% | 99.5% | 99.3% | 98.9% | 99.5% | 99.6% | 99.8% | 96.8% | 99.3% | 99.5% | 99.6% | 99.3% | | 92.4% | 99.8% | 92.3% | 95.9% | 96.0% | 95.8% | 95.8% | 95.3% | 98.2% | 98.2% | 98.9% | 99.1% | 96.3% | 91.9% | 98.9% | 91.9% | 96.0% | 91.7% |
| A/snow goose/Missouri... | 92.1% | 91.9% | 92.1% | 92.3% | 91.9% | 92.4% | 92.2% | 92.4% | 89.8% | 92.3% | 92.3% | 92.3% | 92.1% | 92.4% | | 92.4% | 99.6% | 93.1% | 91.0% | 90.3% | 90.3% | 90.3% | 92.6% | 92.4% | 91.5% | 92.1% | 91.0% | 95.2% | 91.7% | 95.2% | 91.5% | 95.2% |
| A/Prachinburi/6231/2004 | 98.4% | 98.6% | 99.6% | 99.5% | 99.1% | 99.6% | 99.8% | 100% | 97.0% | 99.5% | 99.6% | 99.8% | 99.5% | 99.8% | 92.4% | | 92.3% | 96.1% | 96.1% | 96.0% | 96.0% | 95.4% | 98.4% | 98.4% | 99.1% | 99.3% | 96.5% | 91.9% | 99.1% | 91.9% | 96.1% | 91.7% |
| A/northern pintail/Wash... | 91.9% | 91.7% | 91.9% | 92.1% | 91.7% | 92.2% | 92.0% | 92.3% | 90.0% | 92.1% | 92.1% | 92.1% | 91.9% | 92.3% | 99.6% | 92.3% | | 93.3% | 91.2% | 90.5% | 90.5% | 90.5% | 92.4% | 92.2% | 91.7% | 91.9% | 91.2% | 95.4% | 91.5% | 95.4% | 91.7% | 95.4% |
| A/Japanese white eye/H... | 95.7% | 95.2% | 95.8% | 95.6% | 95.4% | 95.8% | 95.9% | 96.1% | 94.0% | 95.8% | 95.8% | 95.9% | 95.6% | 95.9% | 93.1% | 96.1% | 93.3% | | 94.7% | 94.2% | 94.2% | 93.8% | 96.5% | 96.4% | 95.6% | 95.4% | 95.0% | 93.3% | 95.6% | 93.3% | 96.1% | 93.1% |
| A/Hong_Kong/485/1997 | 95.6% | 96.0% | 95.8% | 95.6% | 95.4% | 95.8% | 95.9% | 96.1% | 94.0% | 96.1% | 95.8% | 96.0% | 95.6% | 96.0% | 91.0% | 96.1% | 91.2% | 94.7% | | 98.6% | 98.6% | 98.2% | 96.0% | 95.8% | 95.6% | 95.4% | 98.2% | 90.8% | 95.4% | 90.8% | 95.2% | 90.7% |
| A/Hong_Kong/483/1997 | 95.4% | 95.4% | 96.0% | 95.4% | 95.2% | 95.6% | 95.8% | 96.0% | 93.8% | 95.6% | 95.6% | 95.8% | 95.4% | 95.8% | 90.3% | 96.0% | 90.5% | 94.2% | 98.6% | | 100% | 97.7% | 95.2% | 95.1% | 95.4% | 95.2% | 97.4% | 90.1% | 95.1% | 90.1% | 94.2% | 90.0% |
| A/Hong_Kong/483/97 | 95.4% | 95.4% | 96.0% | 95.4% | 95.2% | 95.6% | 95.8% | 96.0% | 93.8% | 95.6% | 95.6% | 95.8% | 95.4% | 95.8% | 90.3% | 96.0% | 90.5% | 94.2% | 98.6% | 100% | | 97.7% | 95.2% | 95.1% | 95.4% | 95.2% | 97.4% | 90.1% | 95.1% | 90.1% | 94.2% | 90.0% |
| A/Hong_Kong/481/1997 | 94.9% | 95.1% | 95.1% | 94.9% | 94.7% | 95.1% | 95.2% | 95.4% | 93.3% | 95.4% | 95.1% | 95.3% | 94.9% | 95.3% | 90.3% | 95.4% | 90.5% | 93.8% | 98.2% | 97.7% | 97.7% | | 95.1% | 94.9% | 95.3% | 94.7% | 97.7% | 90.1% | 95.1% | 90.1% | 94.7% | 90.1% |
| A/Hong_Kong/213/2003 | 97.7% | 97.4% | 98.1% | 97.9% | 97.5% | 98.1% | 98.2% | 98.4% | 95.8% | 97.9% | 98.1% | 98.2% | 97.9% | 98.2% | 92.6% | 98.4% | 92.4% | 96.5% | 96.0% | 95.2% | 95.2% | 95.1% | | 100% | 97.5% | 97.7% | 96.5% | 92.3% | 97.5% | 92.3% | 96.7% | 92.1% |
| A/Hong_Kong/212/03 | 97.6% | 97.3% | 98.0% | 97.8% | 97.5% | 98.0% | 98.2% | 98.4% | 96.6% | 97.8% | 98.0% | 98.2% | 97.8% | 98.2% | 92.4% | 98.4% | 92.2% | 96.4% | 95.8% | 95.1% | 95.1% | 94.9% | 100% | | 97.5% | 97.6% | 96.4% | 92.0% | 97.6% | 92.0% | 96.6% | 91.8% |
| A/Hatay/2004 | 97.5% | 97.7% | 98.8% | 98.6% | 98.2% | 98.8% | 98.9% | 99.1% | 96.5% | 98.6% | 98.8% | 98.9% | 98.6% | 98.9% | 91.5% | 99.1% | 91.7% | 95.6% | 95.6% | 95.4% | 95.4% | 95.3% | 97.5% | 97.5% | | 98.4% | 96.0% | 91.4% | 98.6% | 91.4% | 95.6% | 91.2% |
| A/Hanoi/30408/2005 | 98.1% | 97.9% | 98.9% | 98.8% | 99.1% | 98.9% | 99.1% | 99.3% | 96.5% | 98.8% | 98.9% | 99.1% | 98.8% | 99.1% | 92.1% | 99.3% | 91.9% | 95.4% | 95.4% | 95.2% | 95.2% | 94.7% | 97.7% | 97.6% | 98.4% | | 95.8% | 91.5% | 98.4% | 91.5% | 95.8% | 91.4% |
| A/goose/Guangdong/1/... | 96.0% | 95.6% | 96.1% | 96.0% | 95.8% | 96.1% | 96.3% | 96.5% | 94.4% | 96.3% | 96.1% | 96.3% | 96.0% | 96.3% | 91.0% | 96.5% | 91.2% | 95.0% | 98.2% | 97.4% | 97.4% | 97.7% | 96.5% | 96.4% | 96.0% | 95.8% | | 91.0% | 95.8% | 91.0% | 95.1% | 90.8% |
| A/dairy cattle/Texas/24-... | 91.7% | 91.4% | 91.5% | 91.7% | 91.4% | 91.5% | 91.7% | 91.9% | 89.6% | 91.5% | 91.7% | 91.7% | 91.5% | 91.9% | 95.2% | 91.9% | 95.4% | 93.3% | 90.8% | 90.1% | 90.1% | 90.1% | 92.3% | 92.0% | 91.4% | 91.5% | 91.0% | | 91.2% | 100% | 91.4% | 99.5% |
| A/Cambodia/JP52a/2005 | 97.5% | 98.2% | 99.1% | 98.9% | 98.2% | 98.9% | 99.1% | 99.1% | 96.3% | 98.8% | 98.8% | 98.9% | 98.6% | 98.9% | 91.7% | 99.1% | 91.5% | 95.6% | 95.4% | 95.1% | 95.1% | 95.1% | 97.5% | 97.6% | 98.6% | 98.4% | 95.8% | 91.2% | | 91.2% | 95.4% | 91.0% |
| A/Bovine/Ohio/B24OSU... | 91.7% | 91.4% | 91.5% | 91.7% | 91.4% | 91.5% | 91.7% | 91.9% | 89.6% | 91.5% | 91.7% | 91.7% | 91.5% | 91.9% | 95.2% | 91.9% | 95.4% | 93.3% | 90.8% | 90.1% | 90.1% | 90.1% | 92.3% | 92.0% | 91.4% | 91.5% | 91.0% | 100% | 91.2% | | 91.4% | 99.5% |
| A/bar headed goose/Qi... | 97.5% | 95.8% | 95.8% | 95.6% | 95.8% | 95.8% | 95.9% | 96.1% | 94.0% | 96.0% | 95.8% | 96.0% | 95.6% | 96.0% | 91.5% | 96.1% | 91.7% | 96.1% | 95.2% | 94.2% | 94.2% | 94.7% | 96.7% | 96.6% | 95.6% | 95.8% | 95.1% | 91.4% | 95.4% | 91.4% | | 91.4% |
| A/bald eagle/Florida/W2... | 91.5% | 91.2% | 91.4% | 91.5% | 91.2% | 91.3% | 91.5% | 91.7% | 89.4% | 91.4% | 91.5% | 91.6% | 91.4% | 91.7% | 95.2% | 91.7% | 95.4% | 93.1% | 90.7% | 90.0% | 90.0% | 90.1% | 92.1% | 91.8% | 91.2% | 91.4% | 90.8% | 99.5% | 91.0% | 99.5% | 91.4% | |

**Supplemental Figure 1: Phylogenetic relationship between H5 sequences used in this study.** Percentages are expressed as relationship between strains at the amino acid level.

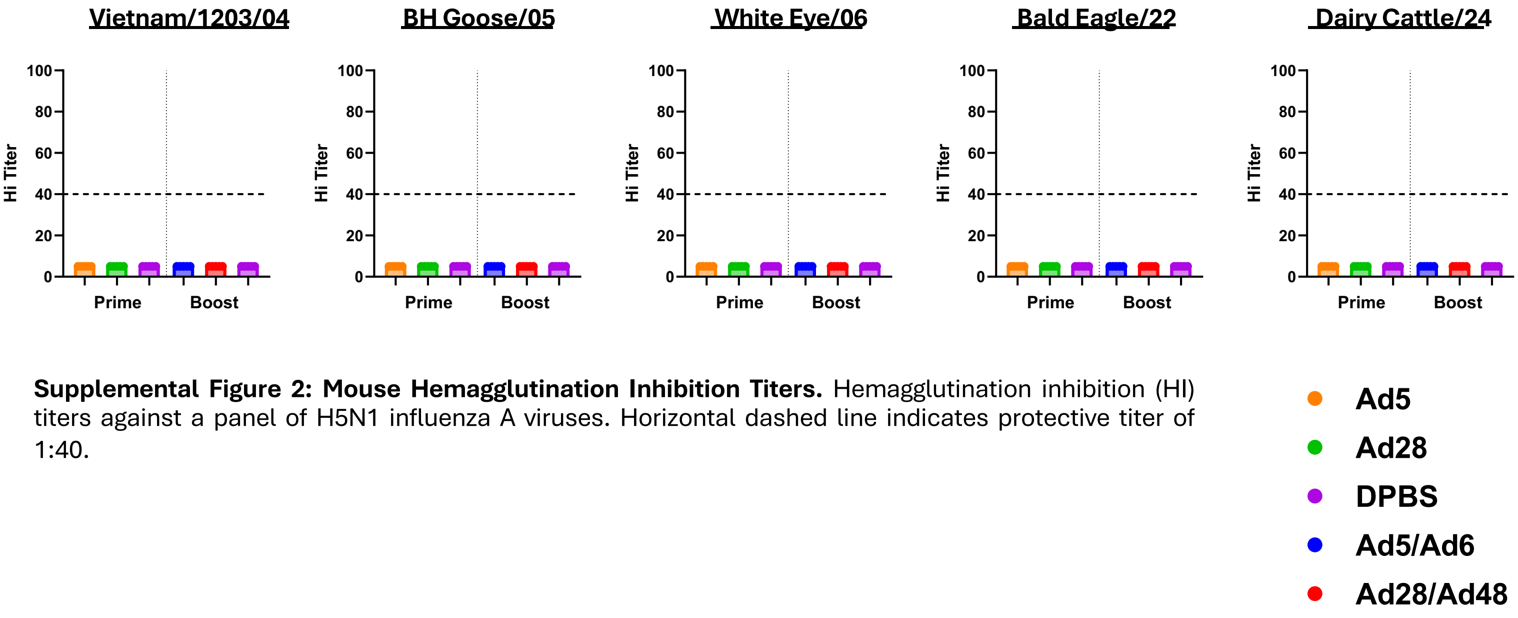


**Supplemental Figure 2: Mouse Hemagglutination Inhibition Titers.** Hemagglutination inhibition (HI) titers against a panel of H5N1 influenza A viruses. Horizontal dashed line indicates protective titer of 1:40.

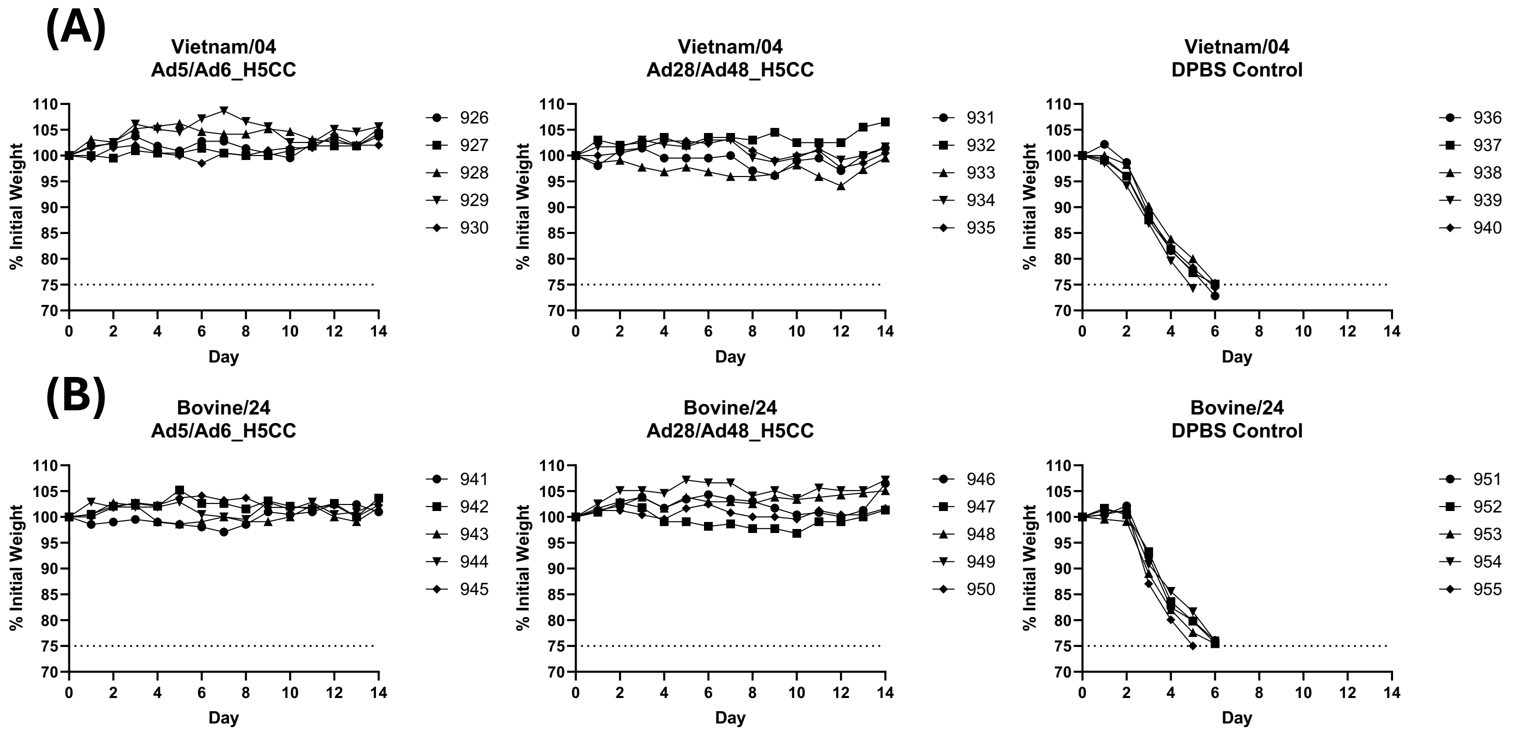


**Supplemental Figure 3: Individual Mouse Weights from IAV Challenge.** (A); A/Vietnam/1203/04 (H5N1) (Vietnam/04). (B); A/Bovine/Ohio/24OSU-439/2024 (H5N1) (Bovine/24). Dashed line indicates the 25% weight loss cut-off used for humane euthanasia.
